# SARS-CoV-2 protein NSP2 enhances microRNA-mediated translational repression

**DOI:** 10.1101/2023.01.01.522328

**Authors:** Parisa Naeli, Xu Zhang, Patric Harris Snell, Susanta Chatterjee, Muhammad Kamran, Reese Jalal Ladak, Nick Orr, Thomas Duchaine, Nahum Sonenberg, Seyed Mehdi Jafarnejad

## Abstract

microRNAs (miRNAs) inhibit mRNA translation initiation by recruiting the GIGYF2/4EHP translation repressor complex to the mRNA 5’ cap structure. Viruses utilise miRNAs to impair the host antiviral immune system and facilitate viral infection by expressing their own miRNAs or co-opting cellular miRNAs. We recently reported that the severe acute respiratory syndrome coronavirus 2 (SARS-CoV-2) encoded non-structural protein 2 (NSP2) interacts with GIGYF2. This interaction is critical for blocking translation of the *Ifn1-b* mRNA that encodes the cytokine Interferon-ß, and thereby impairs the host antiviral immune response. However, it is not known whether NSP2 also affects miRNA-mediated silencing. Here, we demonstrate the pervasive augmentation of the miRNA-mediated translational repression of cellular mRNAs by NSP2. We show that NSP2 interacts with Argonaute 2, the core component of the miRNA-Induced Silencing Complex (miRISC) and enhances the translational repression mediated by natural miRNA binding sites in the 3’ UTR of cellular mRNAs. Our data reveal an additional layer of the complex mechanism by which SARS-CoV-2 and likely other coronaviruses manipulate the host gene expression program through co-opting the host miRNA-mediated silencing machinery.

## Introduction

microRNAs (miRNA) are small, ~22 nucleotides long, non-coding RNAs that modulate the stability and translation efficiency of their target mRNAs. This is mediated by miRNA-Induced Silencing Complex (miRISC), an assembly of a miRNA, Argonaute (AGO), and other proteins, in which miRNA guides the complex to the target mRNA by sequence complementarity[1]. This leads to translational repression followed by deadenylation and decapping of the mRNA, resulting in the exposure of the mRNA to exonuclease-mediated degradation[1–4]. The CCR4-NOT complex plays a key role in coordinating the intricate mechanism of regulation of mRNA translation and decay induced by miRNAs. While miRNA-mediated deadenylation is achieved by the activity of the components of the catalytic subunits of the CCR4-NOT complex (CNOT6/6L and CNOT7/8)[5, 6], translational repression and decapping are engendered through recruitment of several CCR4-NOT binding proteins. We previously showed that recruitment of the mRNA cap-binding eIF4E-homologue protein (4EHP), by CCR4–NOT is critical for the miRNA-mediated translational repression of target mRNAs[7]. 4EHP also forms a translational repressor complex with Grb10-interacting GYF protein 2 (GIGYF2)[8], which represses mRNA translation in CCR4-NOT dependent and independent manners [9]. The GIGYF2/4EHP complex is recruited by a variety of factors including miRNAs [9, 10], the RNA-binding protein Tristetraprolin (TTP) [11], and the stalled ribosome induced Ribosome-associated Quality Control (RQC) mechanism via ZNF598 or EDF1[12] to repress translation.

Viruses use a variety of mechanisms to modulate host gene expression. A common strategy adopted by viruses involves the general shut down of the host mRNAs translation, which allows redirecting the host’s ribosomes toward the viral mRNAs to express the viral proteins[13]. These mechanisms include blocking the cap-dependent translation initiation via sequestering or cleavage of the eukaryotic Initiation factor 4G (eIF4G)[14, 15], binding to and inducing the inhibition or degradation of the poly(A)-binding protein (PABP)[16], and binding to the components of the eIF3 complex[17]. Many RNA viruses bypass the need for cap-dependent translation initiation by utilising an internal ribosome entry site (IRES) in the viral mRNA’s 5’ UTRs to enable translation in a “cap-independent” manner[13]. In addition to shutdown of general cap-dependent translation, viruses also employ “targeted” impairment of the host cell’s homeostasis and proinflammatory responses by changing the expression of miRNAs that target specific host mRNAs[18, 19]. Furthermore, certain viruses express their own miRNAs that target cellular mRNAs[20]. Conversely, host cells also express miRNAs that can interfere with the viral infection by targeting the viral mRNA or silence the antiinflammatory factors[21]. Therefore, the precise regulation of the miRNA-mediated silencing mechanisms is important for the viral infection as well as the host antiviral immune response.

We recently reported that the severe acute respiratory syndrome coronavirus 2 (SARS-CoV-2) encoded non-structural protein 2 (NSP2) functions as a repressor of cellular mRNA translation via direct binding and stabilisation of the GIGYF2/4EHP complex[22]. However, it is not known whether and how NSP2 affects miRNA-mediated silencing, which also utilises the GIGYF2/4EHP complex for translational repression of target mRNAs. Here we provide evidence of a pervasive effect of NSP2 on miRNA-mediated silencing. We show that besides GIGYF2, NSP2 also interacts with the components of miRISC complex and enhances the translational repression of their target mRNAs.

## Results

### NSP2 interacts with the components of miRISC and enhances miRNA-mediated translational repression of target mRNA

To investigate a possible function for the interaction between NSP2 and the GIGYF2/4EHP complex on miRNA-mediated silencing, we first used co-immunoprecipitation (co-IP) to examine the interactions between NSP2 and components of miRISC. We observed that in addition to GIGYF2, NSP2 also interacts with AGO2, the core component of miRISC in HEK293 cells (**Fig. 1A**).

**Figure 1:**
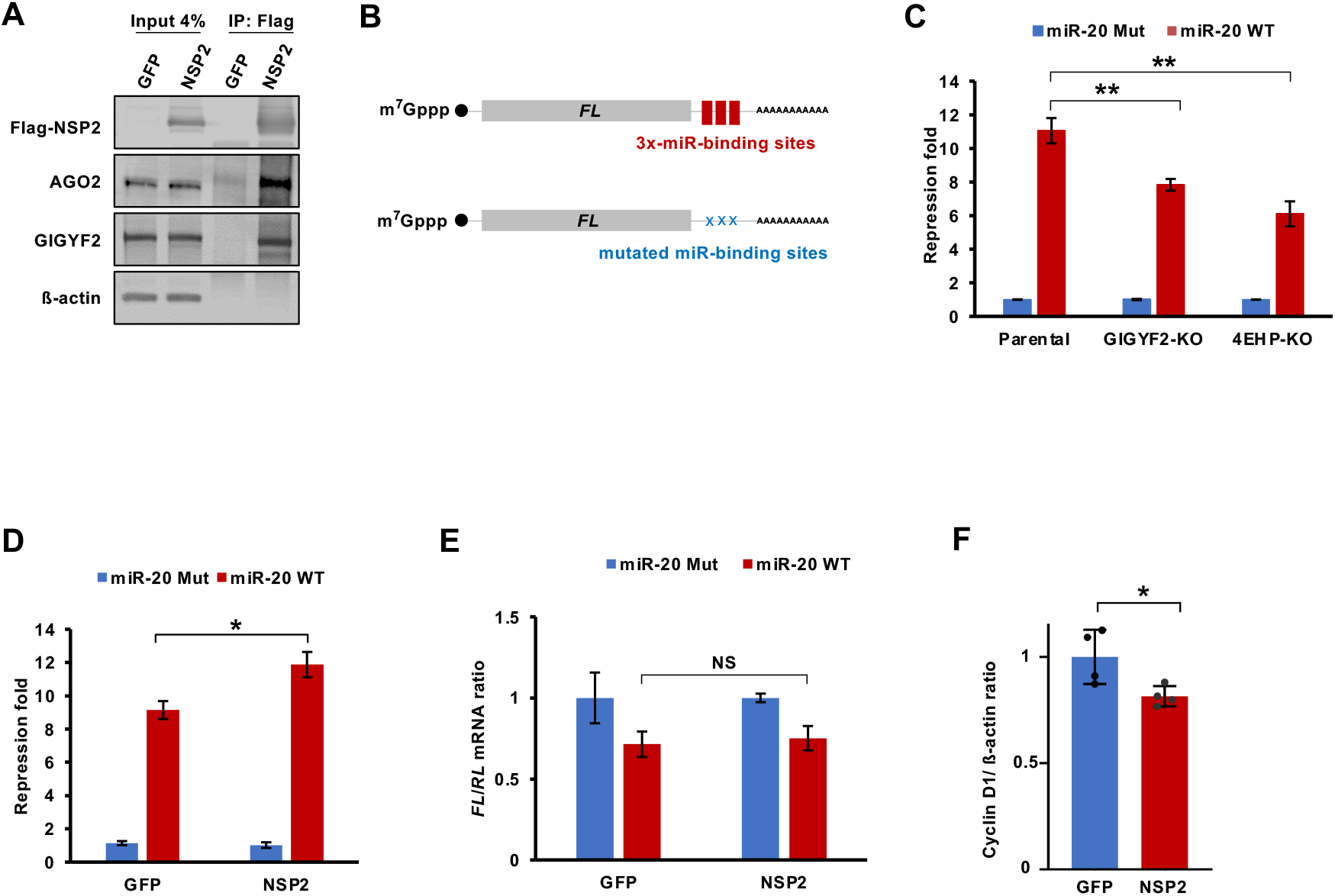
NSP2 interacts with miRISC complex and enhances the miR-20-induced translational silencing. (**A**) HEK293 cells were transfected with Flag-GFP or Flag-NSP2 plasmid. 24 h later, cell lysates were immunoprecipitated using anti-Flag antibody and western blot was performed with the specified antibodies. (**B**) Schematic representation of the *Firefly luciferase (FL)* reporter with 3 miR-20 binding sites (WT) in its 3’ UTR and the control *FL* reporter wherein all 3 miR-20 binding sites are mutated (Mut). (**C**) Parental, GIGYF2-KO, and 4EHP-KO HEK293 cells were co-transfected with WT or Mut *FL*-miR-20 reporter, along with *Renilla luciferase (RL)* as control. Luciferase activity was measured 24 h after transfection. *FL* values were normalized against *RL* levels, and repression fold was calculated for the *FL*-miR-20 WT relative to *FL*-miR-20 Mut level for each population. (**D**) HEK293 cells were transfected with *FL*-miR-20 WT reporter or *FL*-miR-20 Mut reporter (control) and either Flag-GFP or Flag-NSP2. *FL* and *RL* activities were measured 24 h after transfection. *FL* values were normalized against *RL* levels, and repression fold was calculated for the *FL*-miR-20 WT relative to *FL*-miR-20 Mut level for each population. (**E**) RT-qPCR analysis of *FL*-miR-20 WT relative to *FL*-miR-20 Mut expression in Flag-GFP or Flag-NSP2 expressing HEK293 cells, 24 h after transfection. Data are presented as mean ± SD (n=3). NS=non-significant; *p<0.05; **p<0.01 (two-tailed Student’s t-test). (**F**) Quantitation of the western blot analysis of the expression of Cyclin D1 in lysates derived from HEK293 cells that express Flag-NSP2 or Flag-GFP2 as control (see Supp. Fig. 1C for the WB images). Samples were collected from 4 independent biological replicates. ß-actin was used as a loading control. The measured intensity of Cyclin D1 bands were normalised to the corresponding ß-actin bands. Data are presented as mean ± SD (n=4). *p<0.05 (two-tailed Student’s t-test).

We next examined the impact of NSP2 on the repression activity of miRNAs on a reporter mRNA that contains miRNA binding sites. A recent analysis of dysregulated host miRNAs revealed miR-20a as one of the most significantly upregulated miRNAs in SARS-CoV-2 infected cells[23], likely due to its immunosuppressive effects[24, 25]. We used a dual luciferase reporter system in which the *Firefly Luciferase (FL)* mRNA contains 3x miR-20a-binding sites within its 3’ UTR (miR-20 WT; **Fig. 1B**). As a control a similar reporter was used in which all three miRNA binding sites were mutated (miR-20 Mut; **Fig. 1B**). Expression of *Renilla Luciferase (RL)* was used for normalization. We first determined whether the repression of the *FL* reporter by miR-20 is mediated by GIGYF2 and 4EHP by transfecting the parental, GIGYF2-knockout (KO), and 4EHP-KO HEK293 cells with miR-20 WT or miR-20 Mut reporters (**Fig. S1A**). Measurement of the luciferase activity 24 hours after transfection revealed a substantial and significant de-repression of the miR-20 WT reporter in GIGYF2-KO and 4EHP-KO cells compared with the parental cells (8-, 6-, and 11-fold repression respectively; p<0.01; **Fig. 1C & S1A**). This result demonstrates that miR-20a represses the expression of the target mRNA in a GIGYF2- and 4EHP-dependent manner.

To examine the effect of NSP2 on miR-20-induced repression, HEK293 cells were co-transfected with the miR-20 WT or miR-20 Mut reporters and Flag-NSP2 or Flag-GFP as control. We observed that while miR-20 WT was significantly repressed compared with miR-20 Mut in GFP expressing cells, co-expression with NSP2 further augmented miR-20-mediated repression by ~30% (9 and 11.9-fold for GFP and NSP2 expressing cells, respectively; p<0.05; **Fig. 1D & S1B**). Following translational repression, miRNAs induce the degradation of their target mRNAs [3]. To test if NSP2-enhanced miR-20a induced silencing is due to augmented mRNA degradation, we measured the mRNA abundance of the *FL* and *RL* reporters by RT-qPCR. This experiment revealed no significant difference in reporter mRNA expression between GFP and NSP2 expressing cells (p>0.05; **Fig. 1E**). Consistently, we also observed an average 19% reduced expression of Cyclin D1, a natural target of miR-20a[26], at protein level upon overexpression of NSP2 (p<0.05; **Fig. 1F & S1C**), in the absence of a detectable effect on *CCND1* mRNA expression (**Fig. S1D**). These data show that NSP2 significantly enhances the translational repression of target mRNAs induced by miR-20a, without affecting mRNA degradation.

### NSP2-induced translational repression by miRNAs is pervasive

Hitherto, our data have revealed that NSP2 augments miR-20-mediated repression. To investigate whether NSP2 also augments the translational repression by other miRNAs, we used two other abundant miRNAs: let-7a, which is also upregulated in SARS-CoV-2 infected cells[27], and miR-92, the expression of which is not known to be modulated by SARS-CoV-2[23]. Like the miR-20 reporter (**Fig. 1B**), we used the dual luciferase reporter system in which the *FL* mRNA contains either 3x miR-92 binding sites (miR-92 WT) or 3x let-7a binding sites (let-7 WT) within its 3’ UTR. As a control a similar reporter was used in which all three miRNA binding sites were mutated (miR-92 Mut or let-7 Mut). NSP2 overexpression resulted in enhanced repression by miR-92 (6- and 8-fold for GFP and NSP2, respectively; p<0.05; **Fig. 2A & S2A**) and let-7 (2.8- and 3.3-fold for GFP and NSP2, respectively; p<0.01; **Fig. 2B & S2B**) in HEK293 cells. As with miR-20, no significant changes in mRNA levels of miR-92 WT and let-7 WT reporters were detected, demonstrating that NSP2-induced enhanced repression is at the translation level (**Fig. 2C & D**).

**Figure 2:**
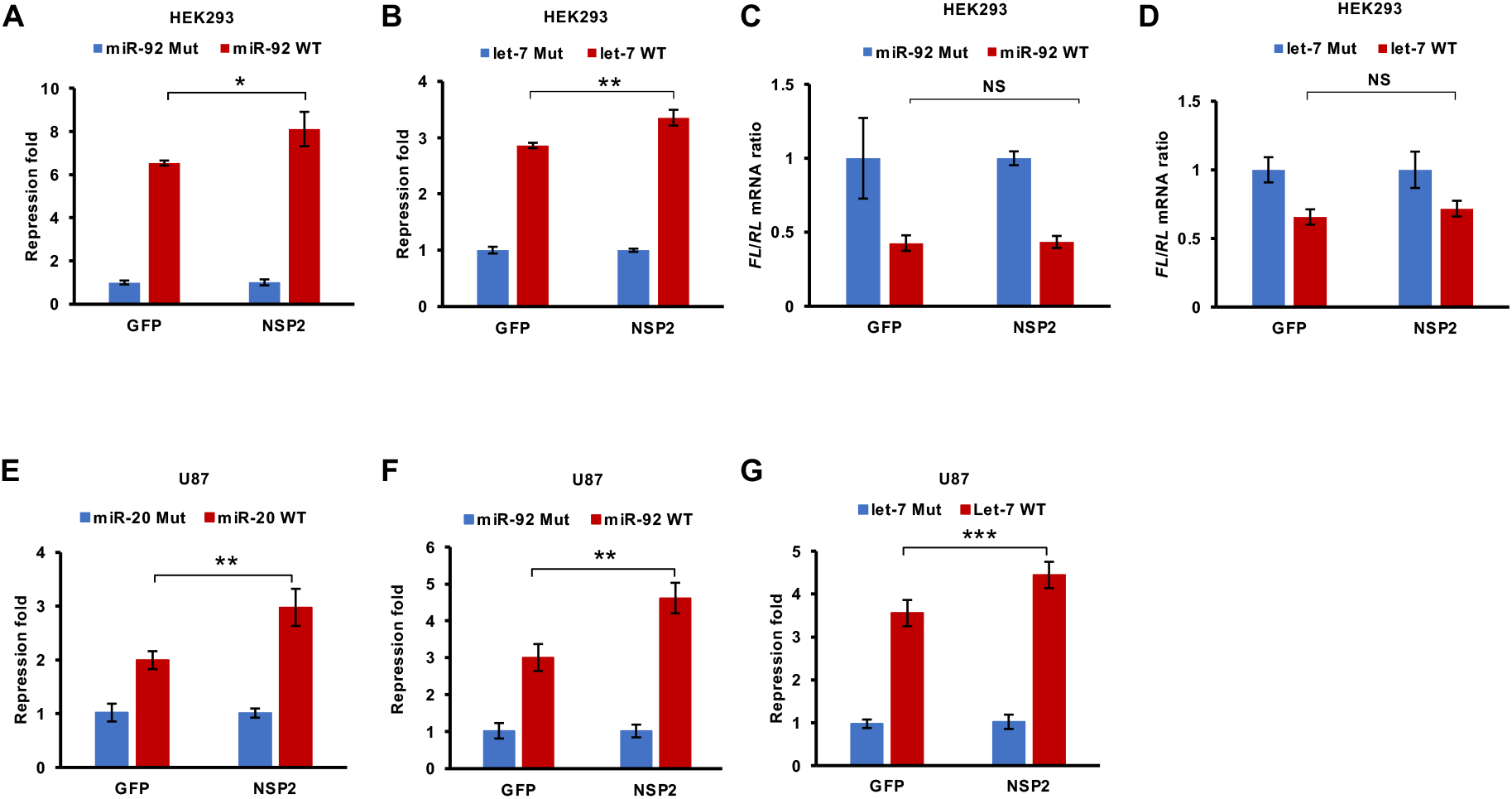
Pervasive effects of NSP2 on miRNA-mediated silencing. (**A** & **B**) HEK293 cells were co-transfected with WT or Mut *FL*-miR-92 reporter (**A**) and WT or Mut *FL*-let-7 reporter (**B**), along with *RL* as control. Luciferase activity was measured 24 h after transfection. *FL* values were normalized against *RL* levels, and repression fold was calculated for the WT relative to the respective Mut levels for each condition. (**C**) RT-qPCR analysis of *FL*-miR-92 WT relative to *FL*-miR-92 Mut expression in Flag-GFP or Flag-NSP2 expressing HEK293 cells, 24 h after transfection. (**D**) RT-qPCR analysis of *FL*-let-7 WT relative to *FL*-let-7 Mut expression in Flag-GFP or Flag-NSP2 expressing HEK293 cells, 24 h after transfection. (**EG**) U87 cells were co-transfected with WT or Mut *FL*-miR-20 ©, *FL*-miR-92 (**F**), or *FL*-let-7 reporter (**G**) along with Flag-NSP2 or Flag-GFP as a control. 24 h later, cells were lysed and luciferases activity was measured. *FL* values were normalized against *RL* levels, and repression fold was calculated for the WT relative to the respective Mut version of the reporter for each condition. Data are presented as mean ± SD (n=3). NS=non-significant; *p<0.05; **p<0.01; ***p<0.001 (two-tailed Student’s t-test).

We next examined whether the observed NSP2-induced enhanced translational repression by miRNAs in HEK293 cells could also be observed in another cell type, the U87 human glioblastoma cell line. Consistent with the results in HEK293 cells, co-expression of NSP2 resulted in significantly enhanced repression by all three tested miRNAs (~50%, ~50%, and ~30% increase for miR-20, miR-92, and let-7 reporters, respectively; **Fig. 2E–G & S2C-E**). Altogether, this data indicates a pervasive role for NSP2 in augmenting miRNA-mediated silencing.

### NSP2 regulates miRNA-mediated silencing of natural 3’ UTR sequences

While bolstering our findings that NSP2 interacts with GIGYF2/4EHP complex, a recent study concluded that this interaction impairs the silencing of target mRNAs induced by let-7 miRNA[28]. We noted that while in our study we used a luciferase reporter with 3x let-7 binding sites (**Fig. 2B & G**), Zou, *et al*. [28] used a reporter with 6x let-7-binding sites. We therefore performed an experiment in which we compared the effects of NSP2 overexpression on reporters with 3x or 6x let-7-binding sites in HEK293T cells (**Fig. 3A & B**). As expected, the repression of the reporter with 3x let-7 binding sites increased by ~60% (2.7- and 4.4-fold in GFP and NSP2 expressing cells respectively; p<0.05; **Fig. 3A**). However, NSP2 overexpression had no significant effect on the 6x let-7 reporter (p>0.05; **Fig. 3B**). We reason that this difference may be due to a saturation of the repression mediated by the highly potent 6x let-7 binding sites (>50-fold repression), which could not be further enhanced by NSP2. In contrast, the let-7 reporter is repressed 10-fold less than the 6x let-7 reporter (<5-fold), which provides a more accurate and physiologically relevant assay for measurement of the impacts of NSP2 on miRNA-mediated silencing.

**Figure 3:**
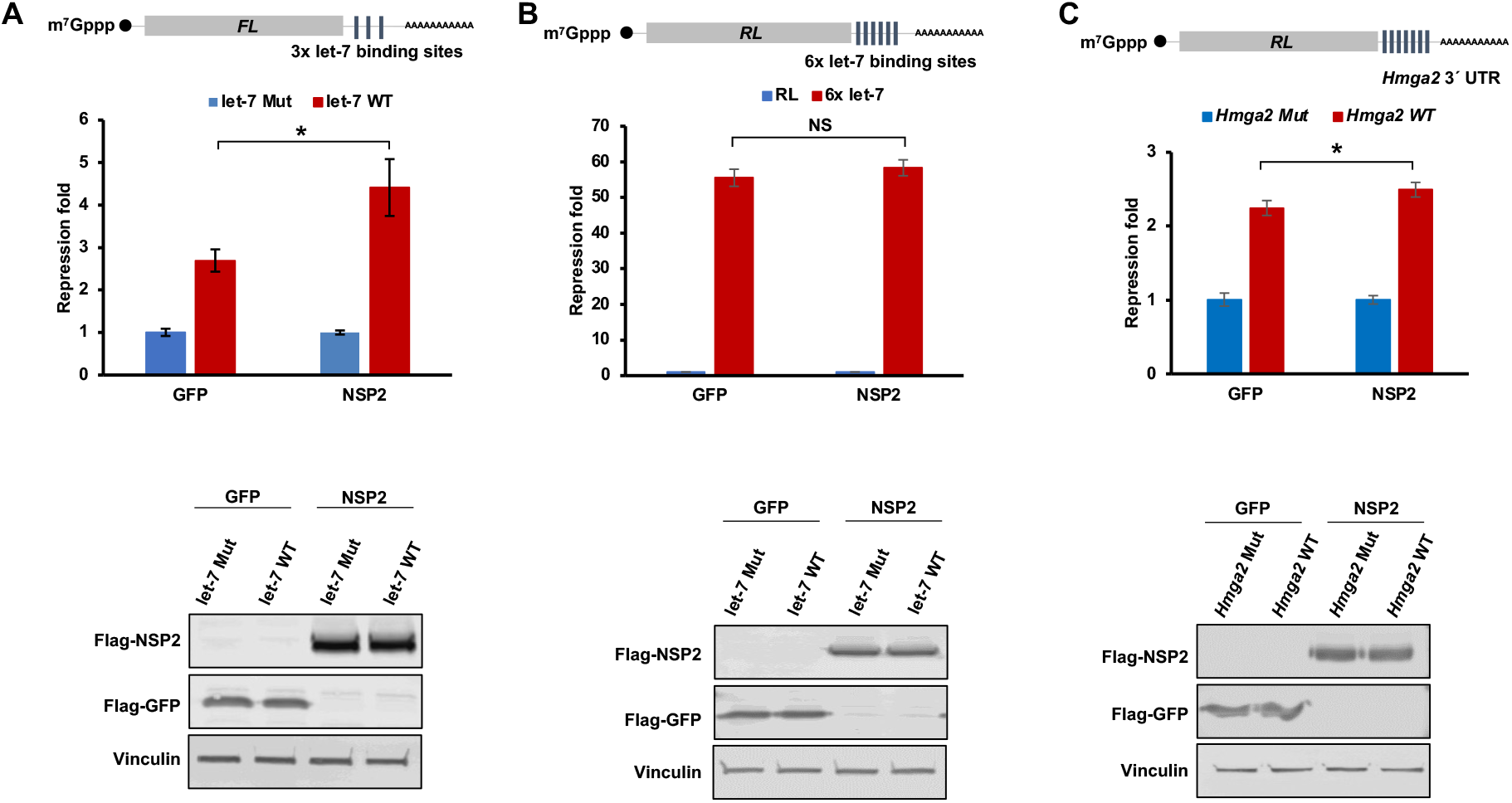
NSP2 enhances the let-7 induced silencing of a reporter with natural 3’ UTR. (**A**) *Top*: HEK293T cells were co-transfected with *FL*-3xlet-7 (WT) reporter or the control version wherein the three let-7 binding sites are mutated (Mut) along with Flag-NSP2 or Flag-GFP as a control. Luciferase activity was measured 24 h after transfection. *FL* values were normalized against *RL* levels, and repression fold was calculated for the WT relative to Mut reporter for each population. *Bottom:* western blot showing the expression of NSP2, GFP, and Vinculin (loading control) corresponding to the top panel. (**B**) *Top*: HEK293T cells were cotransfected with *RL*-6xlet-7 reporter or the control *RL* with no let-7 binding site along with *FL* control and Flag-NSP2 or Flag-GFP as a control. Luciferase activity was measured 24 h after transfection. *RL* values were normalized against *FL* levels, and repression fold was calculated for the *RL*-6xlet-7 relative to the *RL* control reporter for each population. *Bottom:* western blot showing the expression of NSP2, GFP, and Vinculin (loading control) corresponding to the top panel. (**C**) *Top*: HEK293T cells were co-transfected with *RL-Hmga2* 3’ UTR WT or *RL-Hmga2* 3’ UTR Mut reporter, wherein all seven let-7 binding sites are mutated, *FL* plasmid (control), and either Flag-GFP or Flag-NSP2. Luciferase activity was measured after 24 h post-transfection. *RL* values were normalized against *FL* levels, and repression fold was calculated for *RL-Hmga2* 3’ UTR WT relative to the *RL-Hmga2* 3’ UTR Mut reporter for each condition. *Bottom*: western blot showing the expression of NSP2, GFP, and Vinculin (loading control) corresponding to the top panel. Data are presented as mean ± SD (n=3). NS=non-significant; *p<0.05 (two-tailed Student’s t-test).

To further corroborate this conclusion, we tested a reporter fused to the natural (WT) 3’ UTR of the *Hmga2* mRNA, an endogenous target of the let-7 miRNA[29], or a modified version bearing point mutations disrupting all seven let-7 binding sites (**Fig. 3C**) in the presence of NSP2 or a GFP control in HEK293 cells. While the reporter containing the WT *Hmga2* mRNA 3’ UTR was repressed by 2.2-fold compared with the mutated control reporter in GFP expressing cells, this repression was significantly enhanced to 2.5-fold in NSP2 expressing cells (p<0.05; **Fig 3C**). It should be noted that although this natural *Hmga2* mRNA 3’ UTR contains seven let-7 binding site, it induces considerably less repression compared with the 6x let-7 reporter (2-fold and 50-fold repression, respectively). This is likely due to the fact that the natural *Hmga2* 3’ UTR contains additional structures and regulatory elements such as RBP-binding sites [30, 31] that modulate miRNA-mediated silencing [32] but are reflective of physiological repression.

### NSP2 augments AGO2-mediated translational repression

To rule out the possible confounding contributions of changes in miRNA expression upon NSP2 overexpression on our reporter activities, we sought to assess the impact of NSP2 on miRISC activity in a design that is independent of miRNA species. For this, we used the LambdaN (λN):BoxB system, to tether AGO2, which is the minimum required subunit for a functional miRISC complex [33]. A *Renilla luciferase (RL)* encoding five BoxB hairpins in its 3’ UTR (*RL*-5BoxB) was used to tether a λN-AGO2 fusion (**Fig. 4A**). λN-AGO2 repressed the *RL*-5BoxB reporter by 7.7-fold compared with the λN-empty control construct (**Fig. 4A**). Importantly, this repression was reduced to 4.7-fold in GIGYF2-KO and 4.6-fold in 4EHP-KO HEK293 cells (**Fig. 4A & S4A**), which indicates the contribution of a GIGYF2/4EHP-dependent mechanism in AGO2 mediated repression. The prevailing model of miRNA-mediated silencing supports a successive progression from translational repression followed by degradation (decay) of the target mRNA [3]. To specifically dissect the impact on mRNA translation, we used a variation of a *RL*-5BoxB reporter that encodes a self-cleaving hammerhead ribozyme (HhR) at its 3’ end to generate an internalized poly(A) stretch of 114 nucleotides to prevent deadenylation and subsequent degradation of the reporter mRNA (**Fig. 4B**). Tethering of λN-AGO2 resulted in a significant repression of the *RL*-5BoxB-HhR reporter by 3.1-fold (**Fig. 4B**). Importantly, this repression was significantly reduced in GIGYF2-KO and 4EHP-KO cells (to 2.2-fold and 2.5-fold, respectively; **Fig. 4B & S3B**), which indicates the translational repression of the target mRNA by AGO2 is GIGYF2/4EHP-dependent.

**Figure 4:**
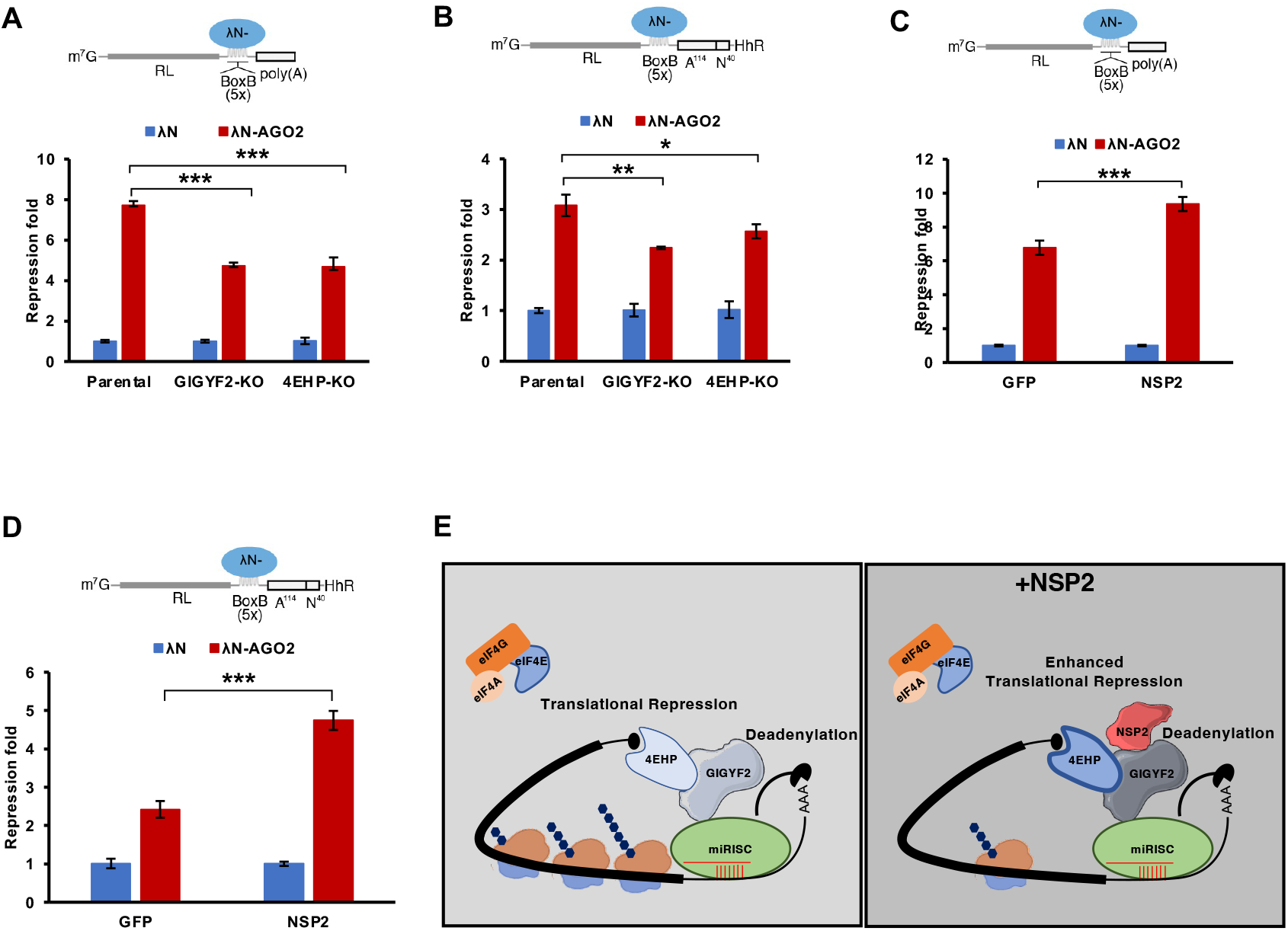
NSP2 augments the miRISC-mediated translational silencing. (**A**) *Top*: schematic representation of the deadenylation permissive *RL*-5BoxB reporter. *Bottom:* Parental, GIGYF2-KO, and 4EHP-KO HEK293 cells were co-transfected with *RL*-5BoxB reporter and either λN-AGO2 or λN-empty along with *FL* plasmid (control). The cells were lysed after 24 h and luciferase activity was measured. *RL* values were normalized against *FL* levels, and repression fold was calculated for the λN-AGO2 relative to λN-empty level for each condition. (**B**) *Top*: schematic representation of the deadenylation-resistant *RL*-5BoxB-HhR reporter. *Bottom:* Parental, GIGYF2-KO, and 4EHP-KO HEK293 cells were co-transfected with *RL*-5BoxB-HhR reporter and either λN-AGO2 or λN-empty along with *FL* plasmid (control). The cells were lysed after 24 h and luciferase activity was measured. *RL* values were normalized against *FL* levels, and repression fold was calculated for the λN-AGO2 relative to λN-empty level for each condition. (**C**) HEK293 cells were co-transfected with the deadenylation permissive *RL*-5BoxB reporter and either λN-AGO2 or λN-empty plasmid, along with *FL* plasmid (control) and either Flag-NSP2 or Flag-GFP. *RL* and *FL* luciferase activity was measured 24 h after transfection and *RL* values were normalized against *FL* levels. Repression fold was calculated for the λN-AGO2 relative to λN-empty level for each condition. (**D**) HEK293 cells were co-transfected with the deadenylation-resistant *RL*-5BoxB-HhR reporter and λN-AGO2 or λN-empty plasmid, along with *FL* plasmid (control) and either Flag-NSP2 or Flag-GFP. *RL* and *FL* luciferase activity was measured 24 h after transfection and *RL* values were normalized against *FL* levels. Repression fold was calculated for the λN-AGO2 relative to λN-empty level for each condition. Data are presented as mean ± SD (n=3). *p<0.05; **p<0.01; ***p<0.001 (two-tailed Student’s t-test). (**E**) Graphic illustration of the mechanism by which NSP2 modulates the cellular miRNA-mediated silencing. The GIGYF2/4EHP complex enables the miRISC-mediated repression of the cap-dependent mRNA translation. Binding of NSP2 to GIGYF2 enhances the interaction between GIGYF2 and 4EHP and the miRNA-mediated translational repression, without affecting the mRNA deadenylation/stability.

Having established this model, we pursued to test the impact of NSP2 on AGO2-mediated silencing. In contrast with the GFP control, overexpression of NSP2 significantly enhanced the λN-AGO2-induced repression of both the degradation-permissive *RL*-5BoxB (**Fig. 4C & S3C**) and the degradation-resistant *RL*-5BoxB-HhR reporters (**Fig. 4D & S3D**). Altogether, these data provide strong evidence that NSP2 enhances mRNAs translational repression by miRISC, independently of the expression or identity of the miRNAs.

## Discussion

Here, we describe a role for the SARS-CoV-2 NSP2 protein in regulation of miRNA-mediated translational repression. We demonstrate that besides GIGYF2, NSP2 also interacts with AGO2 and thereby augments miRNA-mediated translational repression of target mRNAs.

We previously showed that NSP2 facilitates SARS-CoV-2 viral replication by augmenting GIGYF2/4EHP-mediated repression of *Ifnb1* mRNA, which encodes the key cytokine IFN-ß[22]. Notably, *Ifnb1* mRNA contains the binding sites for multiple miRNAs including let-7, miR-34a, miR-26a, and miR-145[34]. Computational analyses identified a substantial number of potential miRNAs encoded by the SARS-CoV-2 genome, many of which were predicted to target mRNAs that encode proteins with important roles in immune-regulatory processes such as NF-κB, JAK/STAT, and TGFß signalling pathways[35]. Further studies empirically identified multiple miRNAs derived from the viral genome, which impaired the host antiviral response by targeting the 3’ UTR of various mRNAs that encode IRF7, IRF9, STAT2[36], and interferon-stimulated genes (e.g. ISG15, MX1, and BATF2)[20, 37, 38]. It is thus likely that enhanced translational repression of targets of both viral and host miRNA through the function of NSP-2 serves to impair a host innate immune response against SARS-CoV-2 infection. However, in principle NSP2 could also enhance the silencing mediated by antiviral miRNAs[39]. To avoid the potential harmful effects of NSP2-induced repression by antiviral miRNAs, the virus can manipulate the expression pattern of antiviral miRNAs [23, 40–43]. Analysis of the expression of 128 human miRNAs with potential to target the SARS-CoV-2 genome revealed their very low expression in lung epithelia[43], which likely allows the virus to avoid the effects of antiviral miRNAs and replicate in these cells.

It is estimated that miRNAs target over 60% of human protein-coding mRNAs[44] and affect important processes including development, cell proliferation, metabolism, and maintenance of homeostasis[45]. Dysregulated miRNA expression and activity has been linked to diseases including cancer and metabolic disorders[46, 47]. Hence, control of miRNA-mediated translational repression by NSP2 during SARS-CoV-2 infection could have significant patho-physiological impacts. Importantly, NSP2, or NSP2-derived peptides that preserve the ability to augment the GIGYF2/4EHP complex could potentially be used to modulate miRNA-mediated silencing, independent of SARS-CoV-2 infection. For instance, global miRNA expression is often downregulated in cancers[48], due to various reasons such as dysregulated expression of miRNA biogenesis factor Dicer[49–51]. It would be interesting to assess the potential effects of enhancement of miRNA-mediated silencing by NSP2 on tumorigenicity of cancers cells with reduced miRNA biogenesis capacity.

Although in the transcriptome-wide scale each miRNA can potentially target hundreds of mRNAs, miRNAs generally have relatively subtle impacts on the stability or translation of individual target mRNAs[52]. In addition, the miRNA-mediated silencing machinery has a limited capacity and is prone to saturation[53, 54]. This limited capacity for miRNA-mediated silencing machinery should be considered when interpreting data generated by transfection of ectopic miRNAs, reporter mRNAs with miRNA-binding sites, or when tethering components of miRISC. Congruent with this notion, NSP2 did not have a significant effect on a reporter mRNA bearing 6x let-7 binding sites, which is repressed by ~50-fold compared with the mutated control (**Fig. 3B**). In contrast, repression of the reporter with 3x let-7 binding sites (**Fig. 3A**) or a reporter with the natural 3’ UTR of the *Hmga2* mRNA, which were repressed by >3-fold was significantly enhanced upon ectopic expression of NSP2 (**Fig. 3C**). This may also explain the discrepancy in the results that we obtained with multiple miRNA reporters (miR-20, miR-92 and let-7) as well as with the tethering of AGO2, compared with the study by Zou *et al*, who used the 6x let-7 reporter or tethering of the GW182 silencing domain that induced >40-fold repression[28].

In addition to mediating the miRNA-induced repression of the cap-dependent mRNA translation, the GIGYF2/4EHP complex can also take part in silencing of target mRNAs triggered by ribosome stalling (Ribosome-associated Quality Control; RQC)[12] or RNA-binding proteins such as TTP[11]. TTP and its paralogues ZFP36L1 and ZFP36L2 have significant roles in cancer[55] and regulation of the immune system[55] through binding and repressing the expression of mRNAs that contain the A/U rich element. A recent study also revealed the biological importance of RQC in neurological disorders [56]. Future studies will assess the impact of NSP2 on other functions of the GIGYF2/4EHP complex, such as TTP and RQC, and the biological pathways they control.

## Materials & Methods

### Cell lines and culture conditions

HEK293T (Thermo Fisher Scientific), and U87 cells (American Type Culture Collection; ATCC) were cultured in DMEM (Dulbecco’s Modified Eagle Medium; Gibco, Cat. # 41965039) supplemented with 10% Foetal Bovine Serum (FBS; Gibco, CAT. # 10270106) and 100 U/ml penicillin, 100 μg/ml streptomycin (Gibco, Cat. # 15070063). 4EHP-KO and GIGYF2-KO HEK293 cells were described previously[19, 22]. Parental, 4EHP-KO, and GIGYF2-KO HEK293 cells were maintained in DMEM supplemented with 10% FBS, 1% penicillin/streptomycin, 100 μg/mL Zeocin (Invitrogen, Cat. # 460509), and 15 μg/mL Blasticidin (Bioshop, Cat. # BLA477). All cells were cultured at 37°C, in a humidified atmosphere with 5% CO_2_ and regularly tested for presence of mycoplasma contamination using mycoplasma detection kit (abm, Cat. # G238).

### Plasmids & cloning

To generate the v5-tagged λN-AGO2 plasmid, the AGO2 CDS was PCR-amplified using pFRT/FLAG/HA-DEST EIF2C2 plasmid (Addgene: Cat. # 19888[57]) as template and Q5 High-Fidelity DNA Polymerase (NEB, Cat. # M0515). AGO2 CDS was subsequently cloned into the pCI-neo-λN-v5 plasmid using the EcoRI and NotI sites. The pmiRGLO plasmid (Promega)-based miR-20, miR-92, and let-7 and the corresponding mutant reporters were described before[58]. The *RL*-6xlet-7 reporter was described previously[59]. The pcDNA3-Flag-GFP and pcDNA3-Flag-NSP2 expressing plasmids were described previously [22].

### Dual luciferase reporter assay

For experiments with miRNAs reporters, 150 x 10^3^ HEK293 or 175 x 10^3^ U87 cells were seeded in 24 well plates and transfected the next day with 100 ng of Flag-NSP2 or Flag-GFP, and 10 ng of wild-type (WT) or mutant (Mut) versions of pmiRGLO-3x-let7-a, pmiRGLO-3xmiR-20a, or pmiRGLO-3xmiR-92 miRNA reporters using Lipofectamine 2000 (Invitrogen, Cat. # 11668019). For experiments with the 6x let-7 reporter, 150 x 10^3^ HEK293T cells were seeded in a 24 well plate and next day were transfected with 100 ng of Flag-NSP2 or Flag-GFP, and 20 ng of *RL*-6xlet-7 and 5 ng of *FL* plasmid as control, using Lipofectamine 2000. The transfection medium was replaced with fresh DMEM + 10% FBS, 6 h after transfection. The cells were lysed 24 h after transfection and luciferase activity was measured with the Dual-Luciferase Reporter Assay kit (Promega Cat. # E1960) according to the manufacturer’s protocol using a FlUOstar plate reader (BMG Labtech). *Firefly Luciferase (FL)* activity was normalized to the *Renilla Luciferase (RL)* activity and the values are shown as repression-fold relative to the control.

For AGO2 tethering, 150 x 10^3^ HEK293 cells were transfected in a 24 well plate with 5 ng of *Firefly Luciferase (FL)* plasmid, 20 ng of *RL*-5BoxB or *RL*-5boxB-A114-N40-HhR, 100 ng of v5-tagged λN-AGO2 or λN-empty and 100 ng of the plasmid encoding Flag-NSP2 or Flag-GFP using Lipofectamine 2000. Cells were lysed 24 h after transfection and luciferase activity was measured with Dual-Luciferase Reporter Assay (Promega Cat. # E1960) according to the manufacturer’s protocol using a FlUOstar plate reader (BMG Labtech). *RL* activity was normalized to the *FL* activity and the repression fold was calculated for λN-AGO2 relative to λN-empty control.

### RNA extraction and RT-qPCR

Total RNA was isolated using TRIzol reagent (Thermo Fisher Scientific, Cat. # 15596026) according to the manufacturer’s instructions. 1 μg purified total RNA was treated with DNase I (Thermo Fisher Scientific, Cat. # EN0521) prior to reverse transcription using SuperScript^™^ III Reverse Transcriptase (Thermo Fisher Scientific, Cat. # 18080085) and random hexamers. The following DNA oligos were used as primers for RCR reactions: pmiRGLO Fluc Forward: ACTTCGAGATGAGCGTTCGG, pmiRGLO Fluc Reverse: CCAACACGGGCATGAAGAAC, pci-neo Fluc Forward: GAGCACGGAAAGACGATGACGG, pci-neo Fluc Reverse: GGCCTTTATGAGGATCTCTCTG, Rluc Forward: ATGGCTTCCAAGGTGTAC, Rluc Reverse: TAGTTGATGAAGGAGTCCA, CCND1 Forward: ACAAACAGATCATCCGCAAACAC, CCND1 Reverse: TGTTGGGGCTCCTCAGGTTC, ß-actin Forward: ACAGAGCCTCGCCTTTGCC, ß-actin Reverse: GATATCATCATCCATGGTGAGCTGG. Quantitative PCR (qPCR) was performed on LightCycler® 480 Instrument II (Roche) using the LightCycler 480 SYBR Green I Master mix (Roche, Cat. # 04887352001), according to the manufacturer’s protocol.

### Western blot

For Western blotting, cells were lysed in RIPA buffer (50 mM TrisHCl pH 7.4, 150 mM NaCl, 2 mM EDTA, 1% NP-40, and 0.1% SDS) supplemented with protease inhibitors (Roche, Cat. # 11836170001). SDS-PAGE gels were used for protein separation followed by transfer onto PVDF membrane (Merck, Cat. # IPFL00010). Blots were blocked with 5% BSA at room temperature for 1 h and incubated with the following primary antibodies overnight at 4°C: mouse anti-v5 (Thermo Fisher Scientific, Cat. # R96025; 1:2000), mouse anti-Flag M2 (Sigma, Cat. # F3165; 1:2000), mouse anti–β-actin (Sigma, Cat. #. A5441; 1:1000), rabbit anti-GIGYF2 (Proteintech, Cat. # 24790-1-AP; 1:1000), rabbit anti-4EHP (Gentex, Cat. # GTX103977; 1:1000), rabbit anti-Vinculin (Cell Signalling, Cat. # 13901S; 1:1000), and rabbit anti-AGO2 (Cell Signalling, Cat # 2897S; 1:1000), mouse anti-Cyclin D1 (Santa Cruz, Cat. # sc-450; 1:1000). The IRDye® 800CW Donkey anti-Rabbit IgG (LI-COR, Cat. # 926-32213) and IRDye® 680RD Donkey anti-Mouse IgG (LI-COR, Cat. # 926-68072) were used as secondary antibodies. All blots were scanned, and images were taken using the Odyssey system (LI-COR, Cat. # ODY-1540). ImageJ software was used for quantification of Cyclin D1 protein expression. The density of Cyclin D1 band was measured and normalized to density of the corresponding ß-actin as loading control. The Cyclin D1 densitometry in NSP2 expressing cell was compared to the GFP expressing cells as control. The uncropped images of all blots are provided in Supplementary Figures 4-8.

### Co-immunoprecipitation (Co-IP)

For Co-IP assays, 4 x 10^6^ HEK293 cells were seeded in 10-cm plates and transfected the next day with 5 μg Flag-GFP or Flag-NSP2 plasmids using Lipofectamine2000 (Thermo Fisher Scientific, Cat. # 11668019), according to manufacturer’s instructions. After 24 h, cells were washed with cold PBS and collected by scraping in the lysis buffer (40 mM HEPES pH 7.5, 120 mM NaCl, 1 mM EDTA, 0.3% CHAPS), supplemented with complete EDTA-free protease inhibitor tablet (Roche, Cat. # 11836170001)]. After 30 min incubation on ice with end-to-end rotation, the lysates were separated from debris by centrifugation at 14,000 g for 15 min at 4°C. The supernatants were pre-cleared by incubation with 50 μl washed and blocked Dynabeads Protein G (Thermo Fisher Scientific, Cat. # 10004D) for 1 h at 4°C. 2 mg pre-cleared lysate was incubated with 3 μg anti-Flag antibody, 60 μl Dynabeads Protein G, and 1 μl RNase I (Invitrogen, Cat. # AM2294) with end-to-end rotation at 4°C overnight. Beads were washed 3×10 min with wash buffer (50 mM HEPES pH 7.5, 150 mM NaCl, 1 mM EDTA, 0.3% CHAPS, and complete EDTA-free protease inhibitor tablet). Protein was eluted in 2x SDS sample buffer for subsequent analysis with western blot.

## Statistical Analyses

Statistical tests were performed using Prism 6 (GraphPad). Error bars represent standard deviation (SD) from the mean of independent replicates. Number of replicates used in each analysis is indicated in the corresponding figure legend. p<0.05 were considered significant.

## Acknowledgment

We are grateful to Prof RJH Davies for critical review and editing of this manuscript and Timothy Winter, Tanvi Nitin Sawant, Brandon Doyle, and Fatema Ali Saif Al Mashaykhi for technical assistance. This work was supported by the Biotechnology and Biological Sciences Research Council (grant # BB/W008165/1; SMJ). MK is supported by a PhD studentship from Brainwaves NI.

## Authors’ contribution

PN performed the majority of the experiments with input from XZ, PHS, SC, MK, and RL. NS, TD, NO, and SMJ contributed to data analysis. PN, NS, and SMJ, conceptualized the study and designed the experiments. SMJ oversaw the study. PN and SMJ wrote the manuscript, which was reviewed, edited, and approved by all authors.

---

**Supplementary Figure 1: Western blot analysis of HEK293 cell lysates; Related to Figure 1.** (**A**) Western blot analysis of lysates from parental, GIGYF2-KO, and 4EHP-KO HEK293 cells transfected with *FL*-miR-20 WT or *FL*-miR-20 Mut with the indicated antibodies. (**B**) Western blot analysis of lysates from HEK293 cells transfected with *FL*-miR-20 WT or *FL-* miR-20 Mut along with Flag-GFP or Flag-NSP2 with the indicated antibodies. (**C**) Western blot analysis of the expression of Cyclin D1 in lysates derived from HEK293 cells that express Flag-NSP2 or Flag-GFP as control (4 independent replicates). ß-actin was used as loading control. (**D**) RT-qPCR analysis of *CCND1* mRNA relative to *ß-actin* expression in Flag-GFP or Flag-NSP2 expressing HEK293 cells, 24 h after transfection. Data are presented as mean ± SD (n=3). NS=non-significant (two-tailed Student’s t-test).

**Supplementary Figure 2: Western blot analysis of HEK293 and U87 cell lysates; Related to Figure 2.** (**A** & **B**) Western blot analysis of lysates from HEK293 cells transfected with *FL-* miR-92 WT or *FL*-miR-92 Mut (**A**) or *FL*-let-7 WT or *FL*-let-7 Mut (**B**) in the presence of Flag-NSP2 or Flag-GFP with the indicated antibodies. (**C-E**) Western blot analysis of lysates from U87 cells transfected with *FL*-miR-20 WT or *FL*-miR-20 Mut (**C**) *FL*-miR-92 WT or *FL*-miR-92 Mut (**D**), *FL*-let-7 WT or *FL*-let-7 Mut (**E**) in the presence of Flag-NSP2 or Flag-GFP with the indicated antibodies.

**Supplementary Figure 3: Western blot analysis of HEK293 cell lysates; Related to Figure 4.** (**A**) Western blot with the indicated antibodies using lysates from the parental, GIGYF2-KO, and 4EHP-KO HEK293 cells transfected with *RL*-5BoxB reporter and λN-AGO2 or λN-empty plasmids. (**B**) Western blot with the indicated antibodies using lysates from the parental, GIGYF2-KO, and 4EHP-KO HEK293 cells transfected with *RL*-5BoxB-HhR reporter and λN-AGO2 or λN-empty plasmids. (**C**) Western blot with the indicated antibodies using lysates from the HEK293 cells transfected with *RL*-5BoxB reporter and λN-AGO2 or λN-empty plasmids, along with Flag-NSP2 or Flag-GFP. (**D**) Western blot with the indicated antibodies using lysates from the HEK293 cells transfected with *RL*-5BoxB-HhR reporter and λN-AGO2 or λN-empty plasmids, along with Flag-NSP2 or Flag-GFP.

**Supplementary Figure 4. Uncropped images of blots used in Figures 1A and Supplementary Figure 1C.**

**Supplementary Figure 5. Uncropped images of blots used in Supplementary Figure 1A, 1B, 2A and 2B.**

**Supplementary Figure 6. Uncropped images of blots used in Supplementary Figure 2C, 2D, 2E and 3A.**

**Supplementary Figure 7. Uncropped images of blots used in Figure 3A, 3B and 3C.**

**Supplementary Figure 8. Uncropped images of blots used in Supplementary Figure 3B, 3C and 3D.**

