## Supplementary Figures 1-8 for "SARS-CoV-2 protein NSP2 enhances microRNA-mediated translational repression"

Suppl. Figure 1

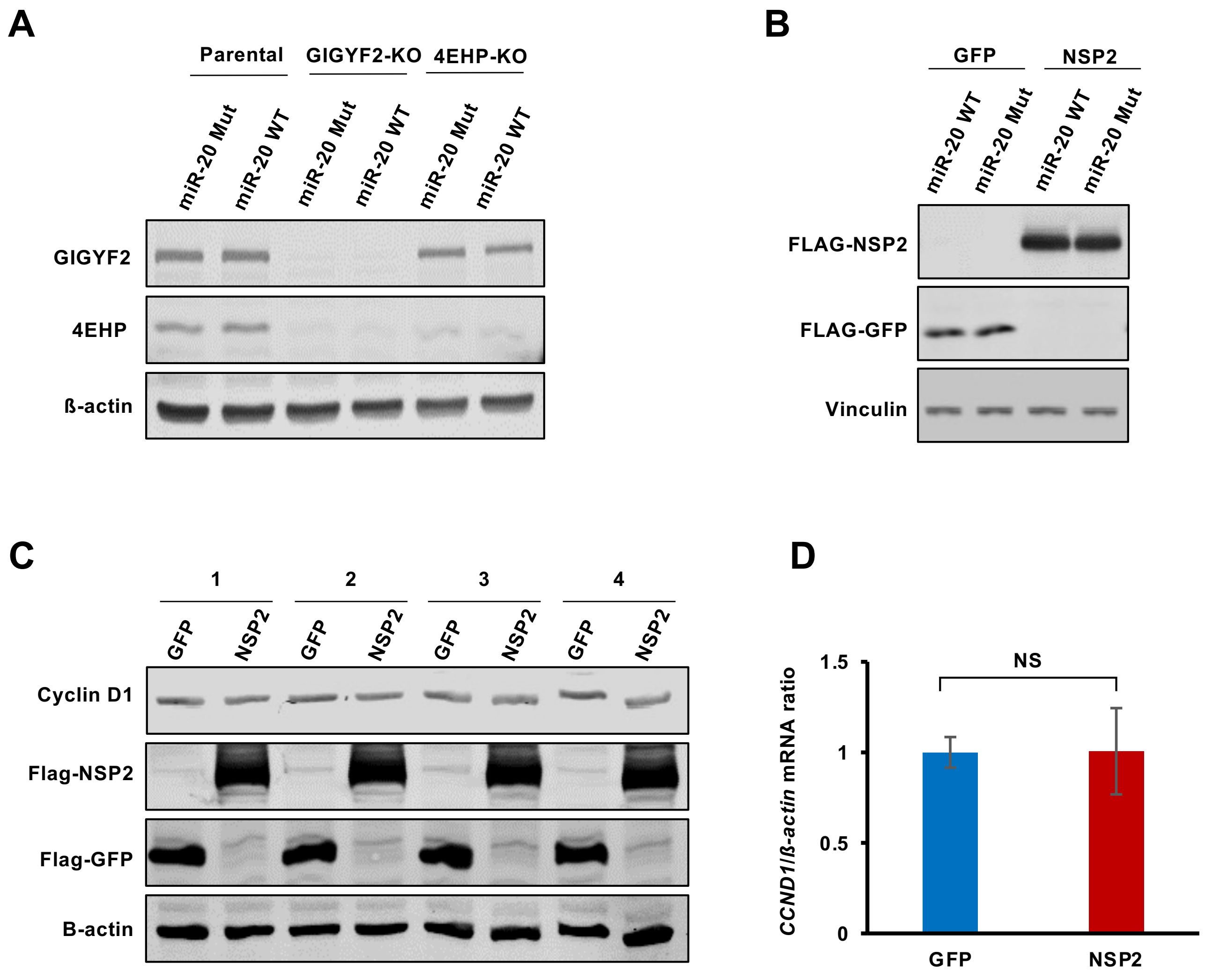

#### Suppl. Figure 2

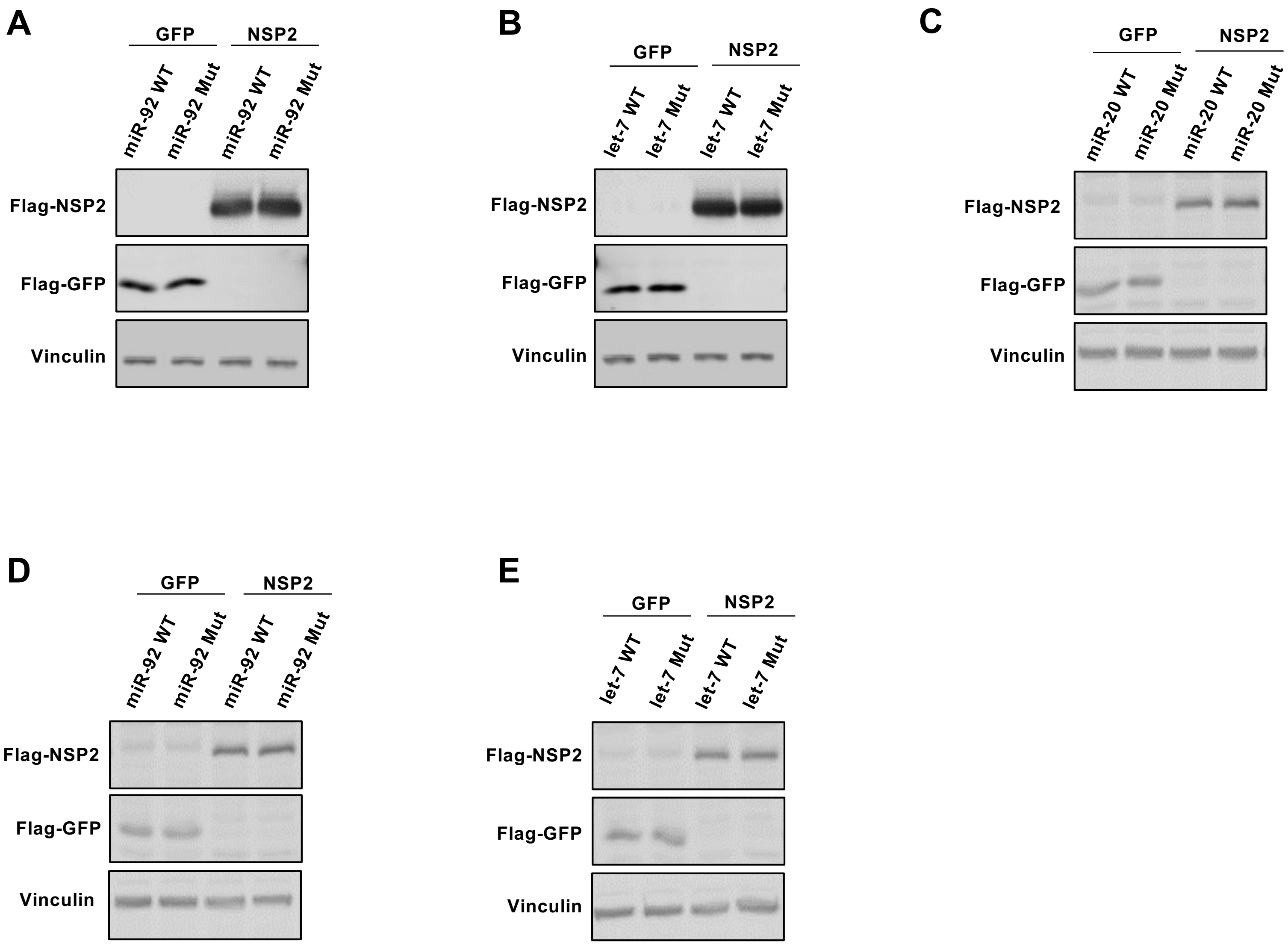

#### Supp. Figure 3

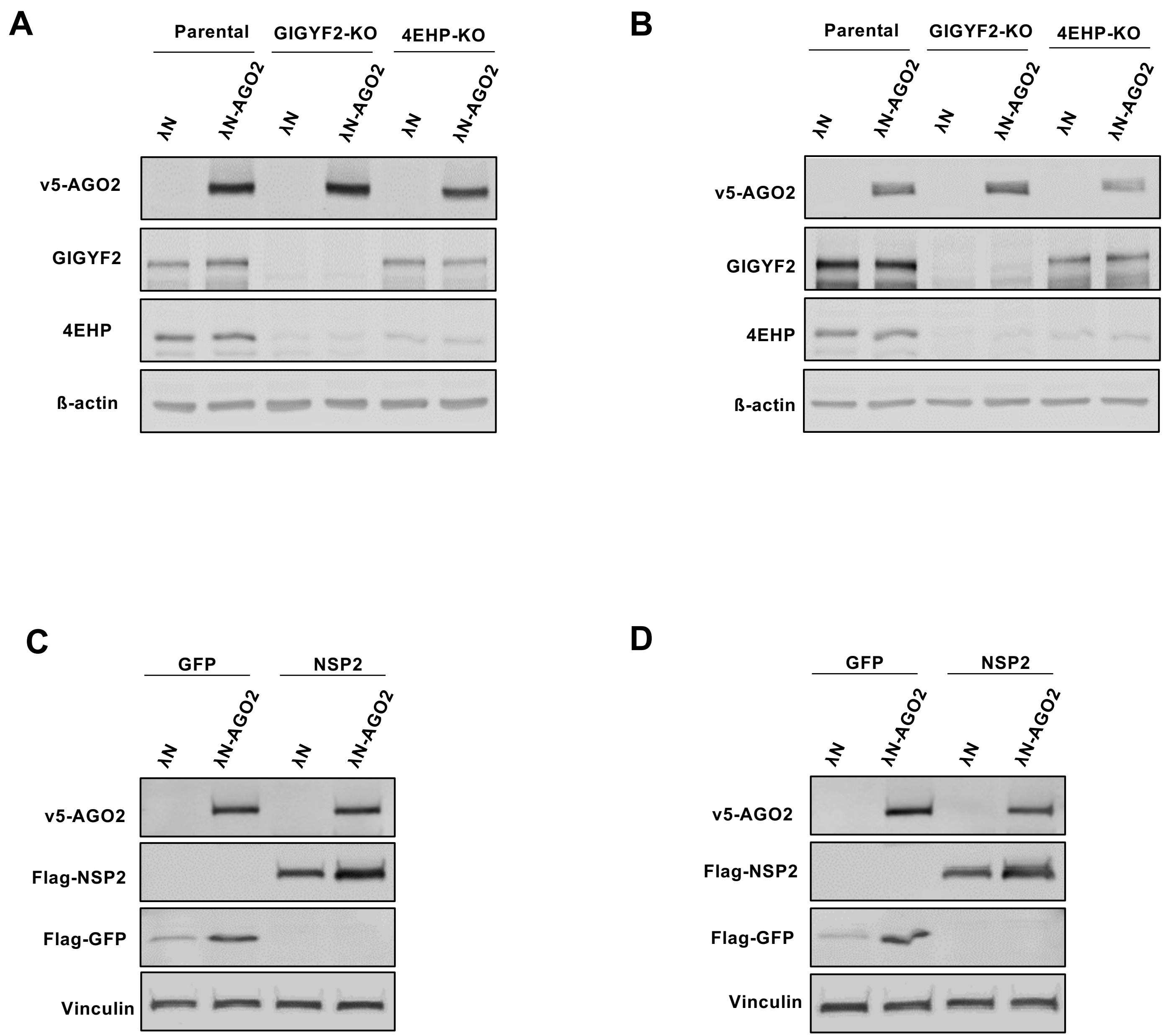

#### Supp. Figure 4

##### Figure 1A

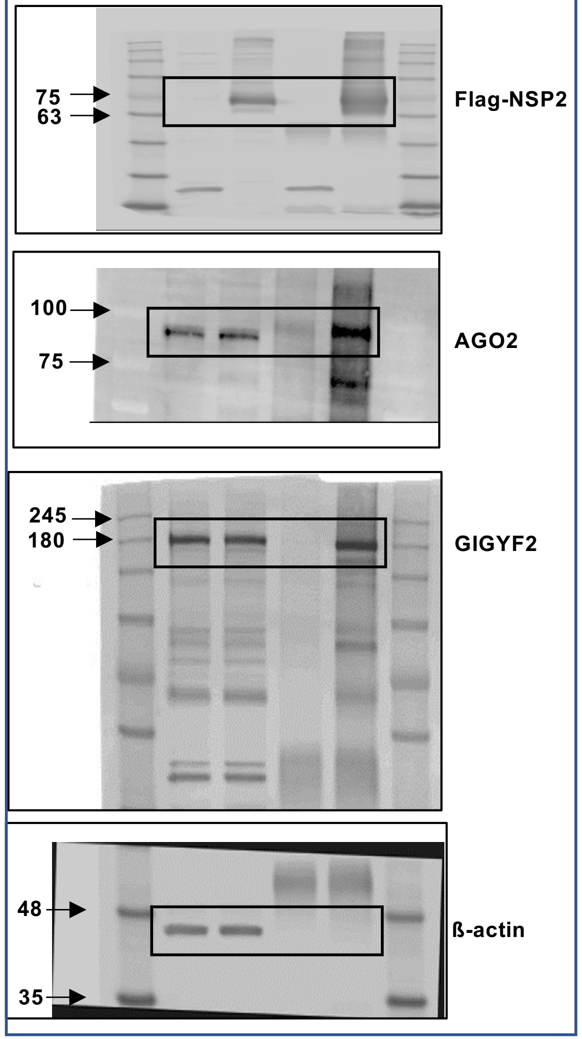


##### Supp. Figure 1C

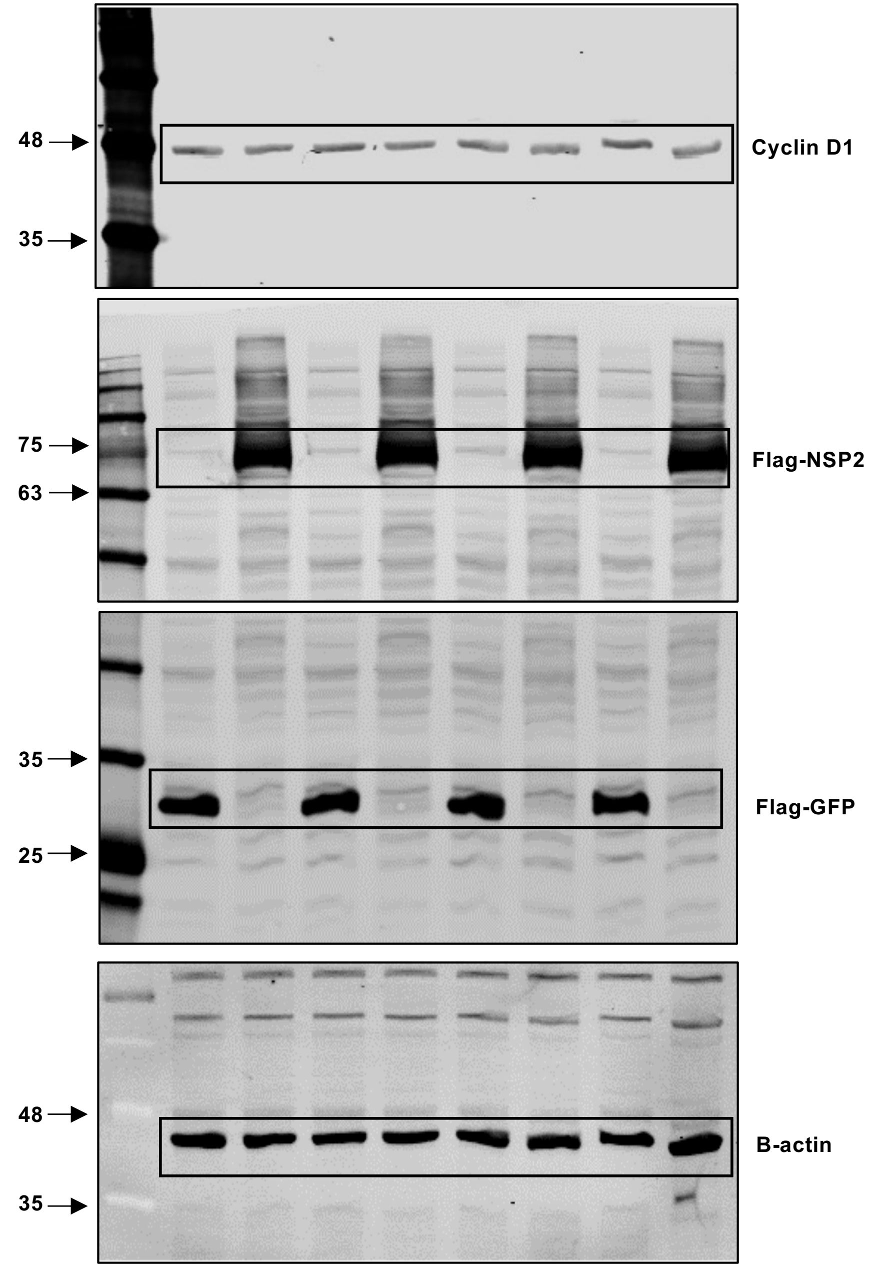

### Supp. Figure 5

#### Supp. Figure 1A

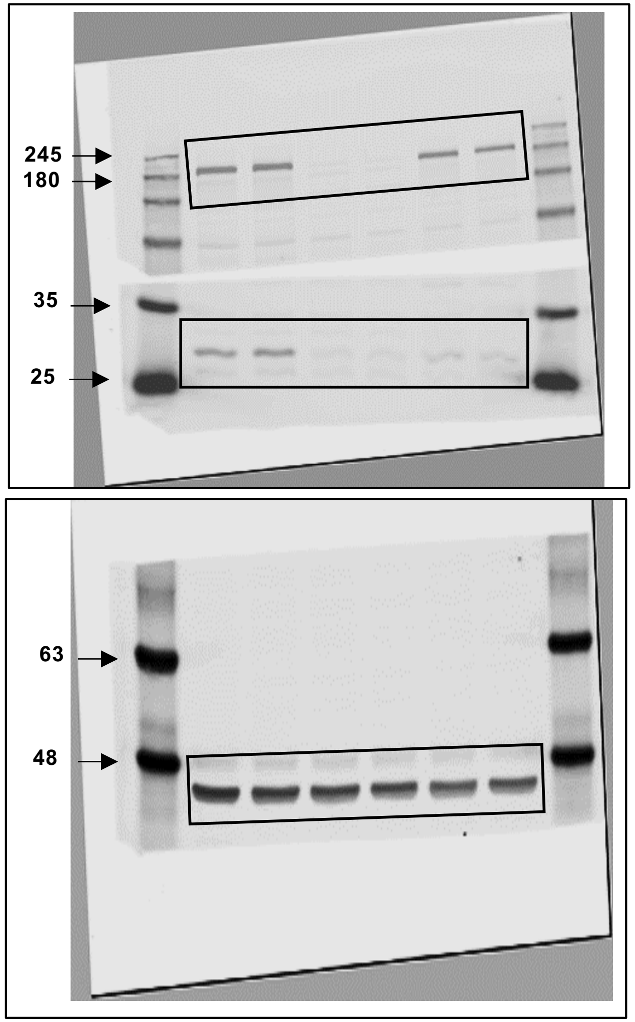


#### Supp. Figure 1B, Supp. Figure 2A & B

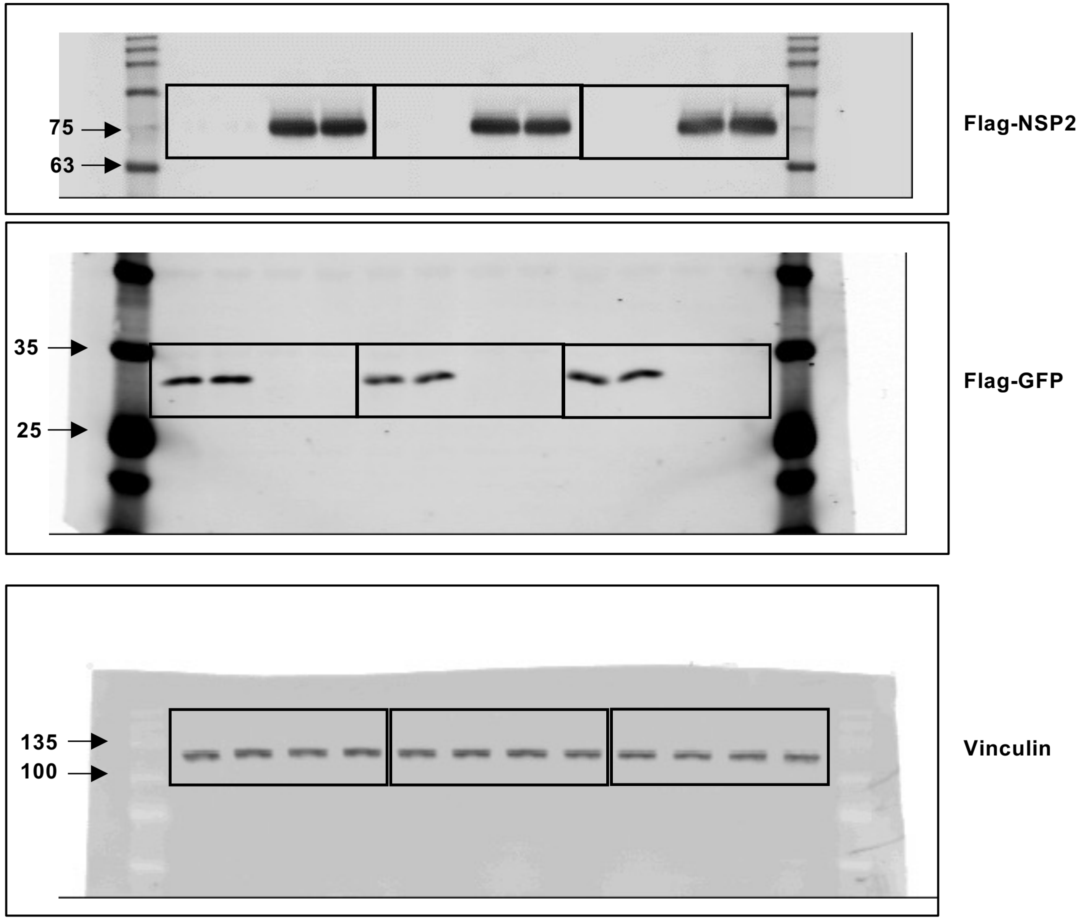

### Supp. Figure 6

#### Supp. Figure 2C, D & E

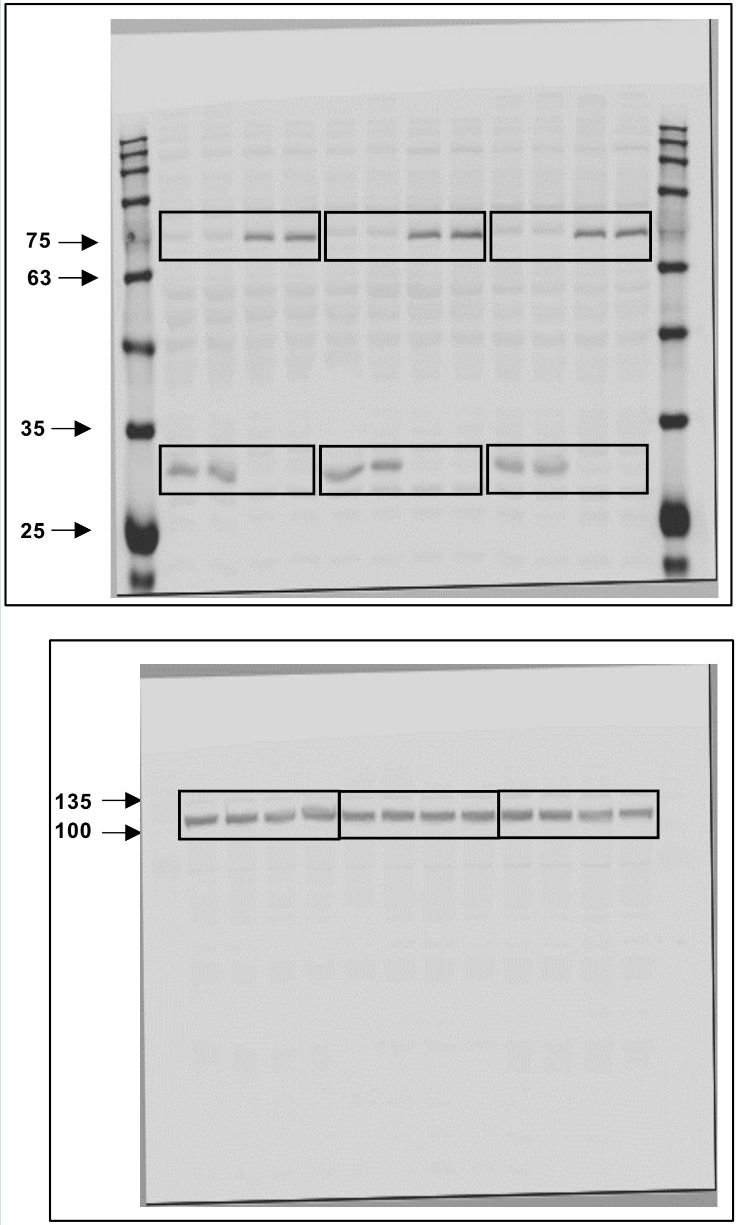


#### Supp. Figure 3A

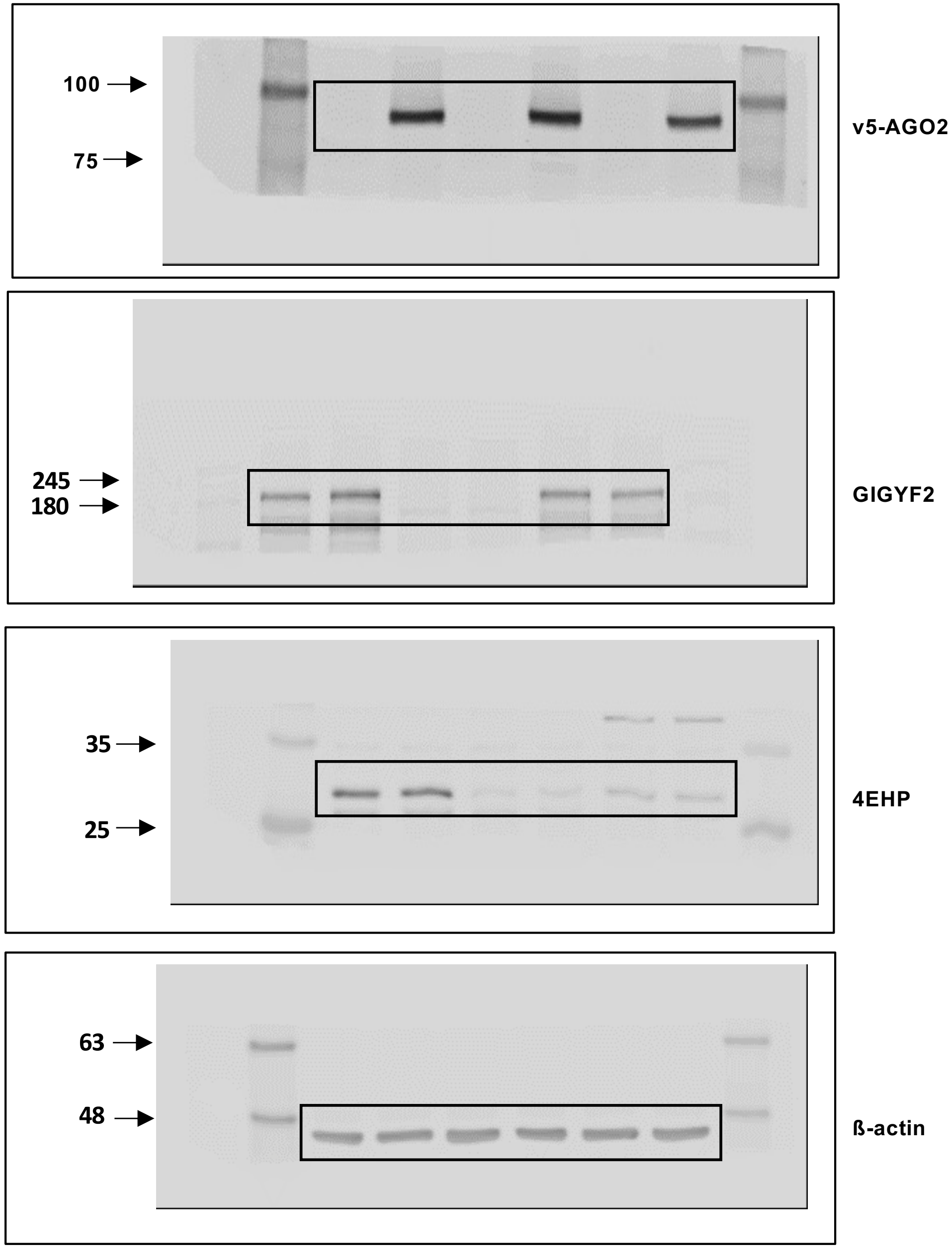

### Supp. Figure 7

#### Figure 3A

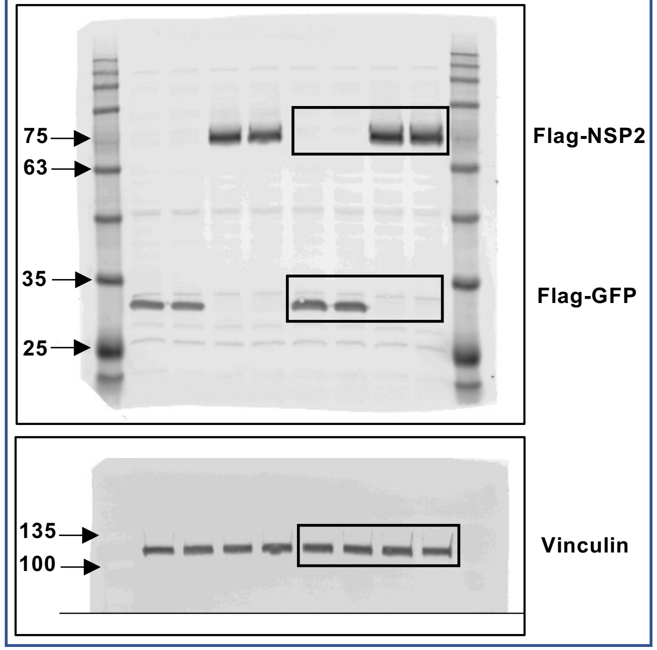


#### Figure 3B

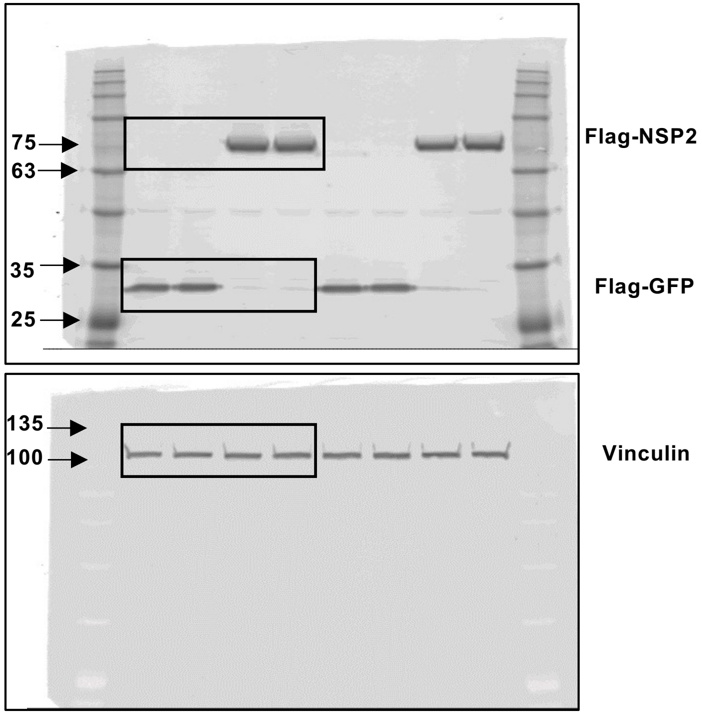


#### Figure 3C

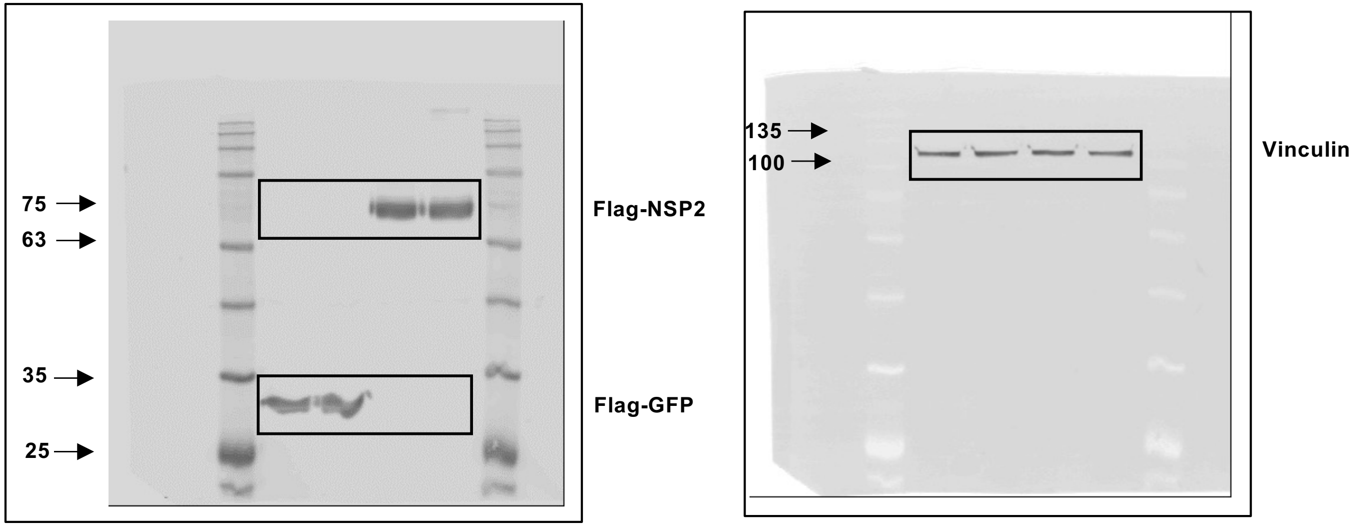

#### Supp. Figure 8

##### Supp. Figure 3B

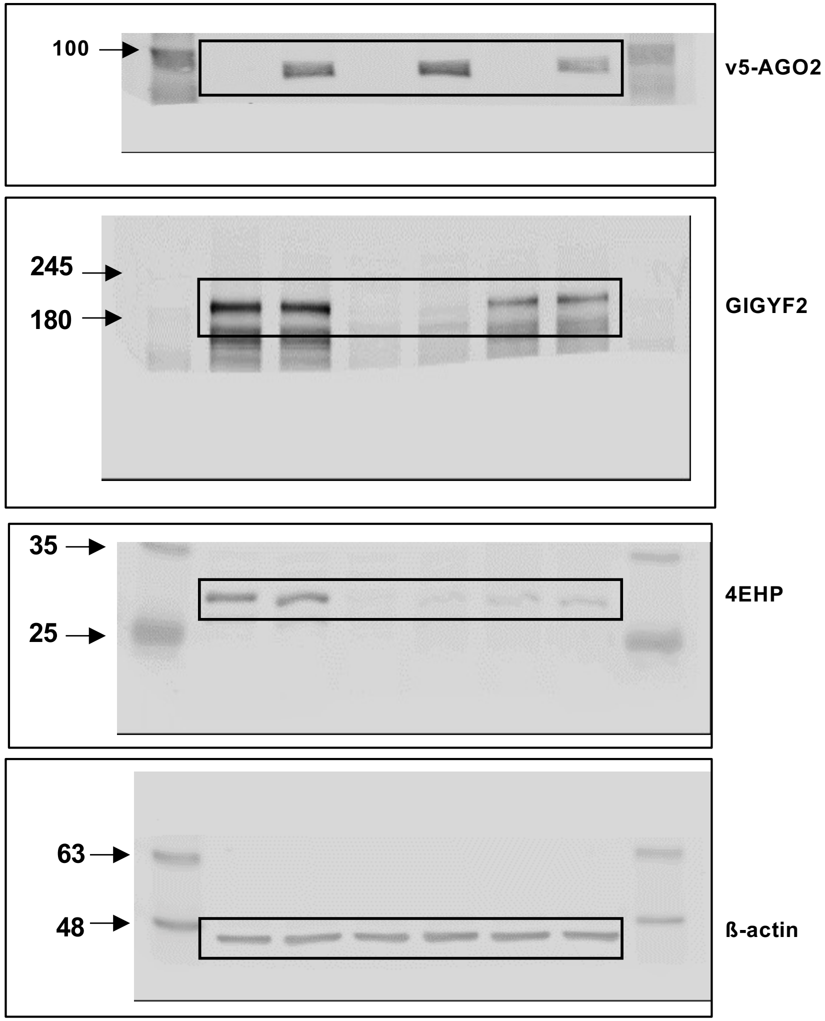


##### Supp. Figure 3C & D

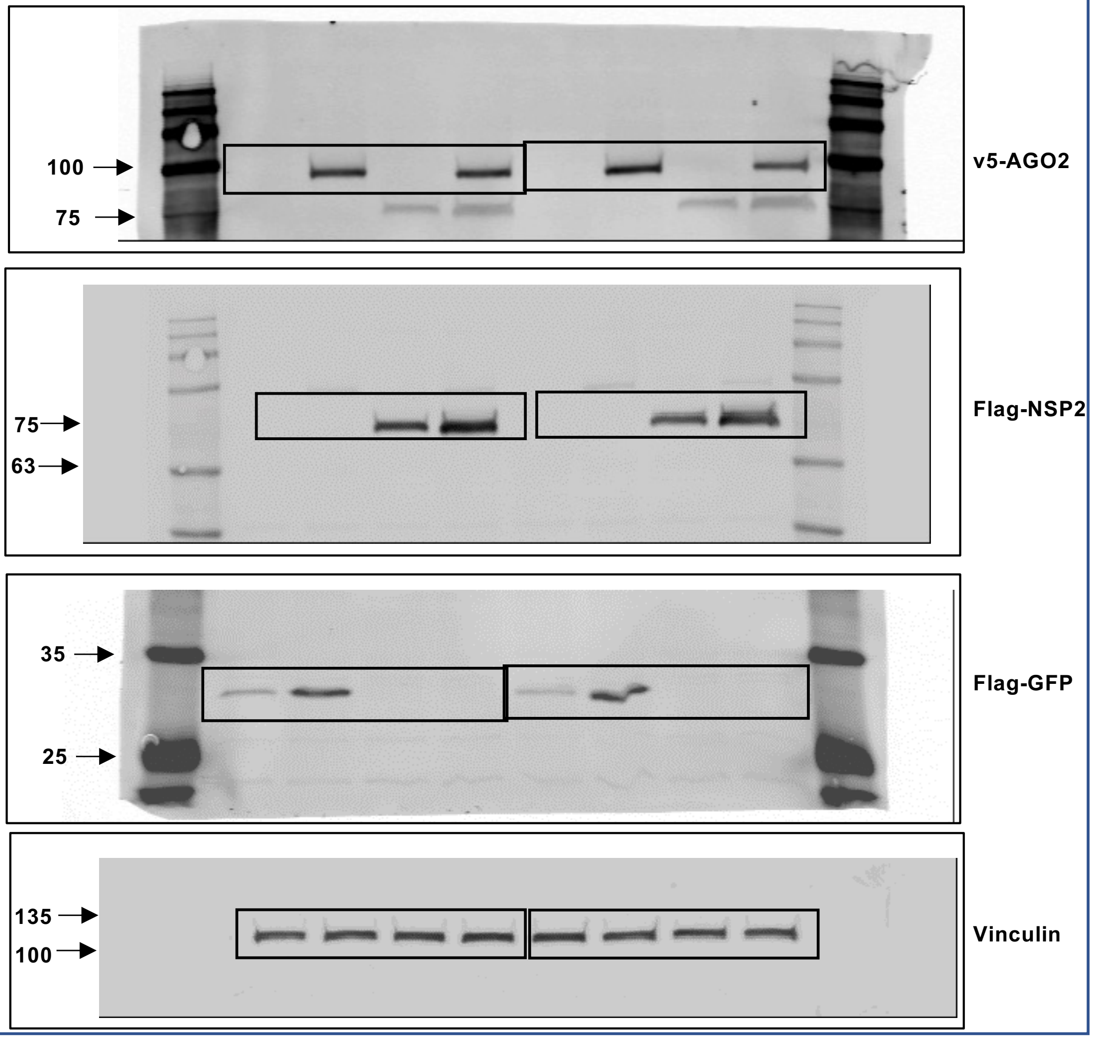
